# Identification of neutralizing human monoclonal antibodies from Italian Covid-19 convalescent patients

**DOI:** 10.1101/2020.05.05.078154

**Authors:** Emanuele Andreano, Emanuele Nicastri, Ida Paciello, Piero Pileri, Noemi Manganaro, Giulia Piccini, Alessandro Manenti, Elisa Pantano, Anna Kabanova, Marco Troisi, Fabiola Vacca, Dario Cardamone, Concetta De Santi, Chiara Agrati, Maria Rosaria Capobianchi, Concetta Castilletti, Arianna Emiliozzi, Massimiliano Fabbiani, Francesca Montagnani, Emanuele Montomoli, Claudia Sala, Giuseppe Ippolito, Rino Rappuoli

## Abstract

In the absence of approved drugs or vaccines, there is a pressing need to develop tools for therapy and prevention of Covid-19. Human monoclonal antibodies have very good probability of being safe and effective tools for therapy and prevention of SARS-CoV-2 infection and disease. Here we describe the screening of PBMCs from seven people who survived Covid-19 infection to isolate human monoclonal antibodies against SARS-CoV-2. Over 1,100 memory B cells were single-cell sorted using the stabilized prefusion form of the spike protein and incubated for two weeks to allow natural production of antibodies. Supernatants from each cell were tested by ELISA for spike protein binding, and positive antibodies were further tested for neutralization of spike binding to receptor(s) on Vero E6 cells and for virus neutralization *in vitro.* From the 1,167 memory B specific for SARS-CoV-2, we recovered 318 B lymphocytes expressing human monoclonals recognizing the spike protein and 74 of these were able to inhibit the binding of the spike protein to the receptor. Finally, 17 mAbs were able to neutralize the virus when assessed for neutralization *in vitro*. Lead candidates to progress into the drug development pipeline will be selected from the panel of neutralizing antibodies identified with the procedure described in this study.

**One Sentence Summary:** Neutralizing human monoclonal antibodies isolated from Covid-19 convalescent patients for therapeutic and prophylactic interventions.

## INTRODUCTION

The impact of the SARS-CoV-2 pandemic, with more than 3.5 million cases, 250,000 deaths and more than 25 million people unemployed in the United States alone, is unprecedented. This first wave of infection is likely to be followed by additional waves in the next few years, until herd immunity, acquired by vaccination or by infection, will constrain the circulation of the virus. It is therefore imperative to develop therapeutic and preventive tools to face the next waves of SARS-CoV-2 infections as soon as possible. Among the many therapeutic options available, human monoclonal antibodies (mAbs) are the ones that can be developed in the shortest period of time. In fact, the extensive clinical experience with the safety of more than 50 commercial mAbs approved to treat cancer, inflammatory and autoimmune disorders provides high confidence on their safety. These advantages, combined with the urgency of the SARS-CoV-2 pandemic, support and justify an accelerated regulatory pathway. In addition, the long industrial experience in developing and manufacturing mAbs decreases the risks usually associated with the technical development of investigational products. Finally, the incredible technical progress in the field allows to shorten the conventional timelines and go from discovery to proof of concept trials in 5-6 months (*1*). Indeed, in the case of Ebola, mAbs were the first therapeutic intervention recommended by the World Health Organization (WHO) and they were developed faster than vaccines or other drugs (*2*).

The SARS-CoV-2 spike glycoprotein (S-protein) has a pivotal role in viral pathogenesis and it is considered the main target to elicit potent neutralizing antibodies and the focus for the development of therapeutic and prophylactic tools against this virus (*3, 4*). Indeed, SARS-CoV-2 entry into host cells is mediated by the interaction between S-protein and the human angiotensin converting enzyme 2 (ACE2) (*3, 5*). The S-protein is a trimeric class I viral fusion protein which exists in a metastable prefusion conformation and in a stable postfusion state. Each S-protein monomer is composed of two distinct regions, the S1 and S2 subunits. Structural rearrangement occurs when the receptor binding domain (RBD) present in the S1 subunit binds to the host cell membrane. This interaction destabilizes the prefusion state of the S-protein triggering the transition into the postfusion conformation which in turn results in the ingress of the virus particle into the host cell (*6*). Single-cell RNA-seq analyses to evaluate the expression levels of ACE2 in different human organs have shown that SARS-CoV-2, through the S-protein, can invade human cells in different major physiological systems including the respiratory, cardiovascular, digestive and urinary systems, thus enhancing the possibility of spreading and infection (*7*).

To identify potent mAbs against SARS-CoV-2 we isolated over a 1,100 S-protein specific-memory B cells derived from seven Covid-19 convalescent donors. As the S-protein RBD domain is mainly exposed when this glycoprotein is in its prefusion state (*6*), we screened naturally produced mAbs against either the S1/S2 subunits and the S-protein trimer stabilized in its prefusion conformation (*6*). This strategy allows us to identify mAbs able to recognize linear epitopes as well as highly neutralizing trimer specific regions on the S-protein surface. The potent neutralizing effect of trimer specific mAbs has already been shown for other pathogens including the respiratory syncytial virus (RSV) (*8, 9*).

In this paper we report the identification of a panel of 318 mAbs from which we plan to select lead candidates for clinical development.

## RESULTS

### SARS-CoV2 induces a strong antibody response in patients contracting infection

Patients recovered from SARS-CoV-2 infection were enrolled in two ongoing clinical studies based in Rome, Italy (National Institute for Infectious Diseases, IRCCS, Lazzaro Spallanzani) and Siena, Italy (Azienda Ospedaliera Universitaria Senese). We firstly examined whether these patients had anti-SARS-CoV-2 S-protein antibodies. Plasma samples were evaluated by enzyme linked immunosorbent assay (ELISA) to assess the polyclonal response to the S-protein trimer, for their ability to neutralize the binding of the spike protein to Vero E6 cells (neutralization of binding or NOB assay) and for their potency in neutralizing the cytopathic effect caused by SARS-CoV-2 infection *in vitro.* Results shown in Table 1 and Figure 1 show that, among the seven donors included in this study, six were able to produce high titers of SARS-CoV-2 S-protein specific antibodies and in particular donors R-042, R-122 and R-188 showed the highest virus neutralizing titers. Only one patient (R-276) mounted low anti-spike polyclonal response (Fig. 1A-B). Interestingly, despite no statistically significant correlation was observed when Pearson correlation analysis was performed, it is possible to observe a trend of correlation between S-protein binding, NOB titer and neutralization titer, suggesting that an abundant response against the S-protein trimer in its prefusion conformation may be indicative of immunity against SARS-CoV-2 (Fig. 1C-D). A bigger dataset may be needed to support this observation.

**Fig. 1.**
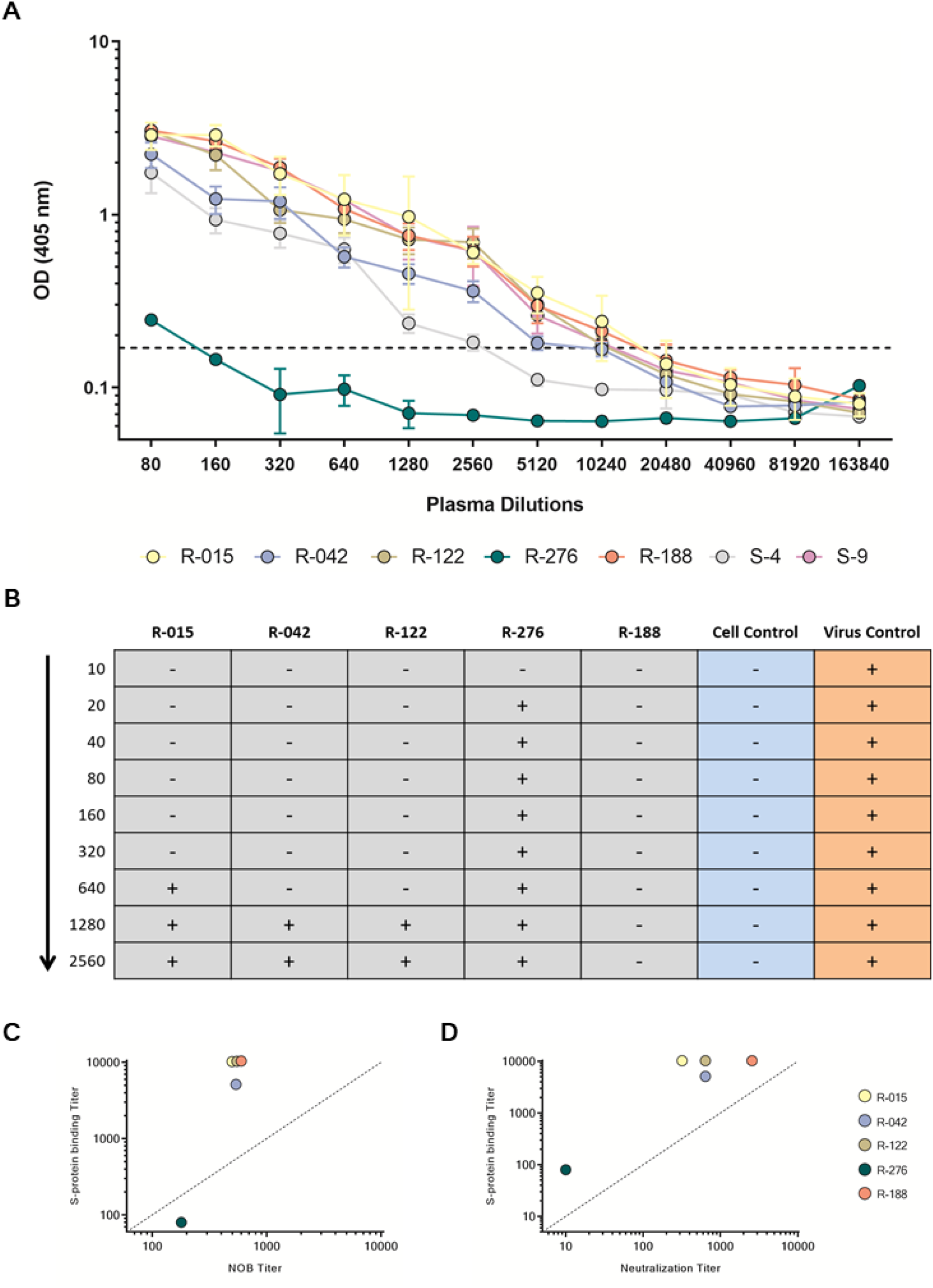
S-protein binding and neutralization titration of SARS-CoV-2 convalescent donors plasma. (A) Plasma samples were two-fold diluted starting at 1:80 to test their ability to bind the S-protein trimer in its prefusion state by ELISA. Results were considered as positive when the OD_405_ value was at least two times higher than the blank. (B) Plasma samples were two-fold diluted starting at 1:10 to test their ability to neutralize SARS-CoV-2 *in vitro.* Results were considered as positive when no cytopathic effect (-) was observed on Vero E6 cells. (C) The graph shows on the Y axis the Log_10_ S-protein binding titer and on the X axis the Log_10_ NOB titer of plasma collected from Covid-19 convalescent patients. Donors R-015 and R-122 are not visible in the graph as their data overlap with those of donor R-188. (D) The graph shows on the Y axis the Log_10_ S-protein binding titer and on the X axis the Log_10_ neutralization titer of plasma collected from Covid-19 convalescent patients.

**Table 1.**
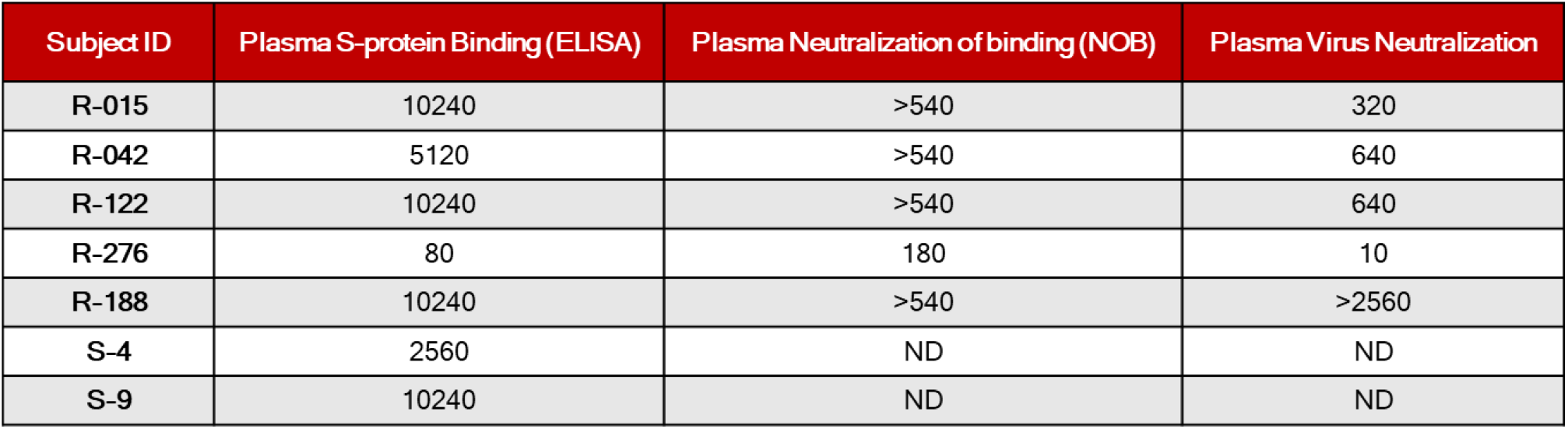
SARS-CoV-2 convalescent donors plasma analyses. Plasma S-protein binding titers for each subject were measured by ELISA assays. Neutralization activity was detected by NOB and by neutralization of SARS-CoV-2 infection of Vero cells; ND = Not Done.

| Subject ID | Plasma S-protein Binding (ELISA) | Plasma Neutralization of binding (NOB) | Plasma Virus Neutralization |
| --- | --- | --- | --- |
| R-015 | 10240 | >540 | 320 |
| R-042 | 5120 | >540 | 640 |
| R-122 | 10240 | >540 | 640 |
| R-276 | 80 | 180 | 10 |
| R-188 | 10240 | >540 | >2560 |
| S-4 | 2560 | ND | ND |
| S-9 | 10240 | ND | ND |

### Isolation of naturally induced S-protein specific antibodies from SARS-CoV-2 convalescent patients

To retrieve mAbs specific for SARS-CoV-2 S-protein, peripheral blood mononuclear cells (PBMCs) from the seven convalescent patients enrolled in this study were collected and stained with fluorescent labeled S-protein trimer to identify antigen specific memory B cells (MBCs). The gating strategy described in Fig. 2 was used to single cell sort into 384-well plates IgG^+^ and IgA^+^ MBCs binding to the SARS-CoV-2 S-protein trimer in its prefusion conformation. A total of 1,167 S-protein-binding MBCs were successfully retrieved with frequencies ranging from 0,17% to 1,41% (Table 2). Following the sorting procedure, S-protein^+^ MBCs were incubated over a layer of 3T3-CD40L feeder cells in the presence of IL-2 and IL-21 stimuli for two weeks to allow natural production of immunoglobulins (*10*). Subsequently, MBC supernatants containing IgG or IgA were tested for their ability to bind either the SARS-CoV-2 S-protein S1 + S2 subunits (Fig. 3A) or trimer in its prefusion conformation (Fig. 3B) by enzyme linked immunosorbent assay (ELISA). A panel of 318 mAbs specific for the SARS-CoV-2 S-protein were identified showing a broad range of signal intensities (Table 2 and Fig. 3).

**Fig. 2.**
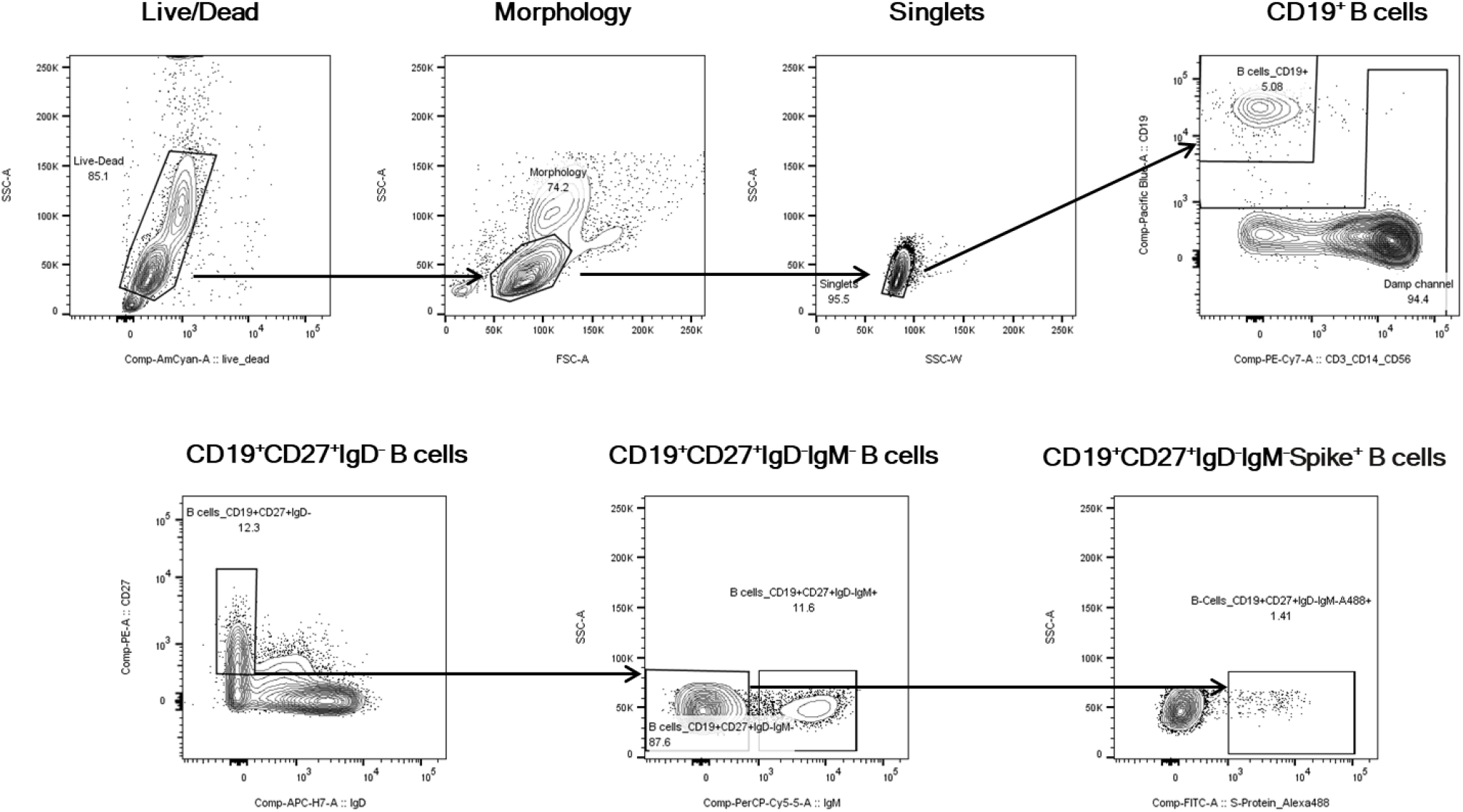
Gating strategy for S-protein specific MBC single cell sorting. Starting from top left to the right panel, the gating strategy shows: Live/Dead; Morphology; Singlets; CD19^+^ B cells; CD19^+^CD27^+^IgD^-^; CD19^+^CD27^+^IgD^-^IgM^-^; CD19^+^CD27^+^IgD^-^IgM^-^S-protein^+^B cells.

**Fig. 3.**
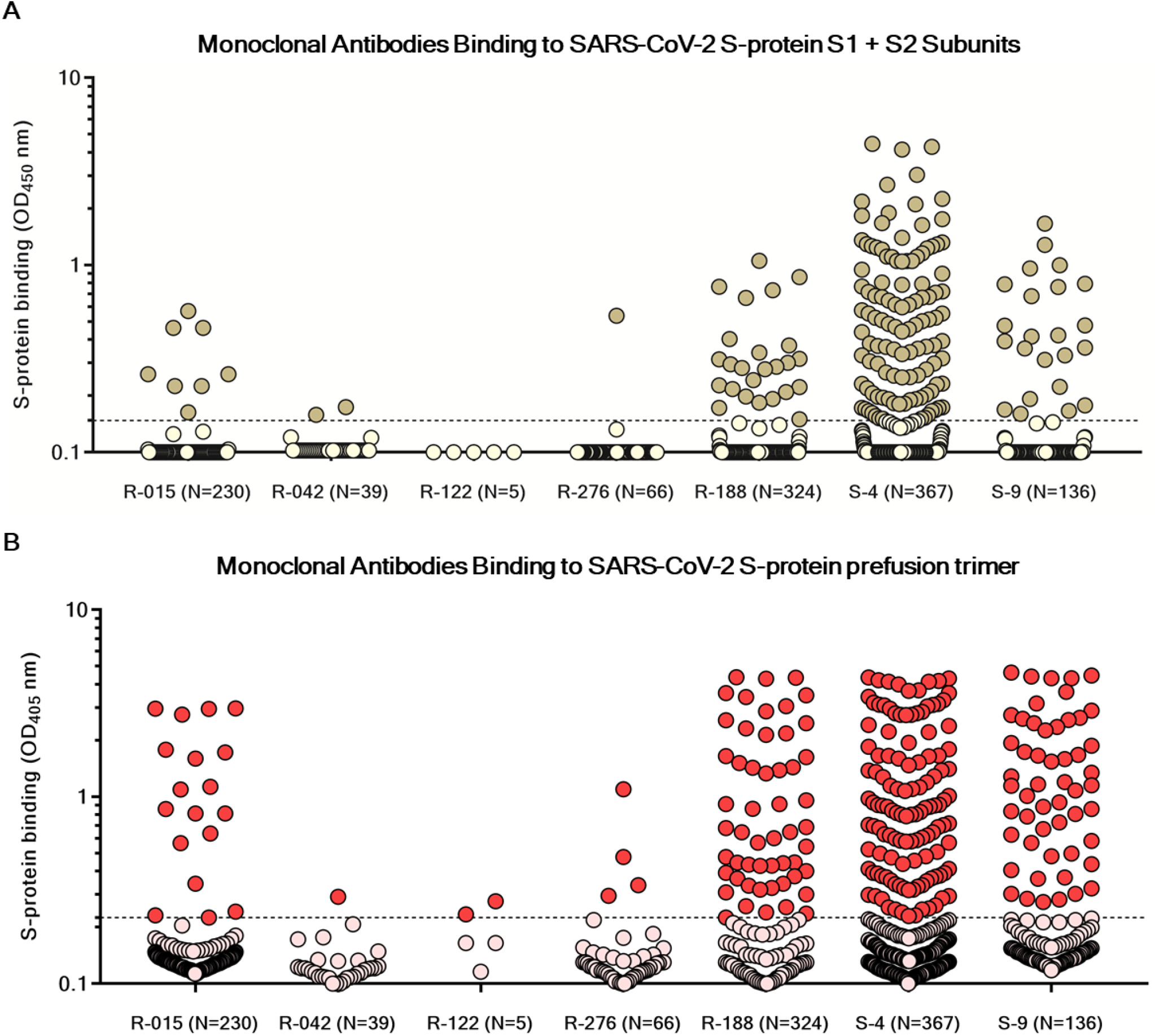
Identification of SARS-CoV-2 S-protein specific mAbs isolated from convalescent donors. (A) The graph shows supernatants tested for binding to the SARS-CoV-2 S-protein S1 + S2 subunits. Threshold of positivity has been set as two times the value of the blank (dotted line) Darker dots represent mAbs which bind to the S1 + S2 while light yellow dots represent mAbs which do not bind. The total number (N) of single cell sorted B cell supernatants screened for binding is also shown for each donor. (B) The graph shows supernatants tested for binding to the SARS-CoV-2 S-protein stabilized in its prefusion conformation. Threshold of positivity has been set as two times the value of the blank (dotted line). Red dots represent mAbs which bind to the S-protein while pink dots represent mAbs which do not bind. The total number (N) of single cell sorted B cell supernatants screened for binding is also shown for each donor.

**Table 2.**
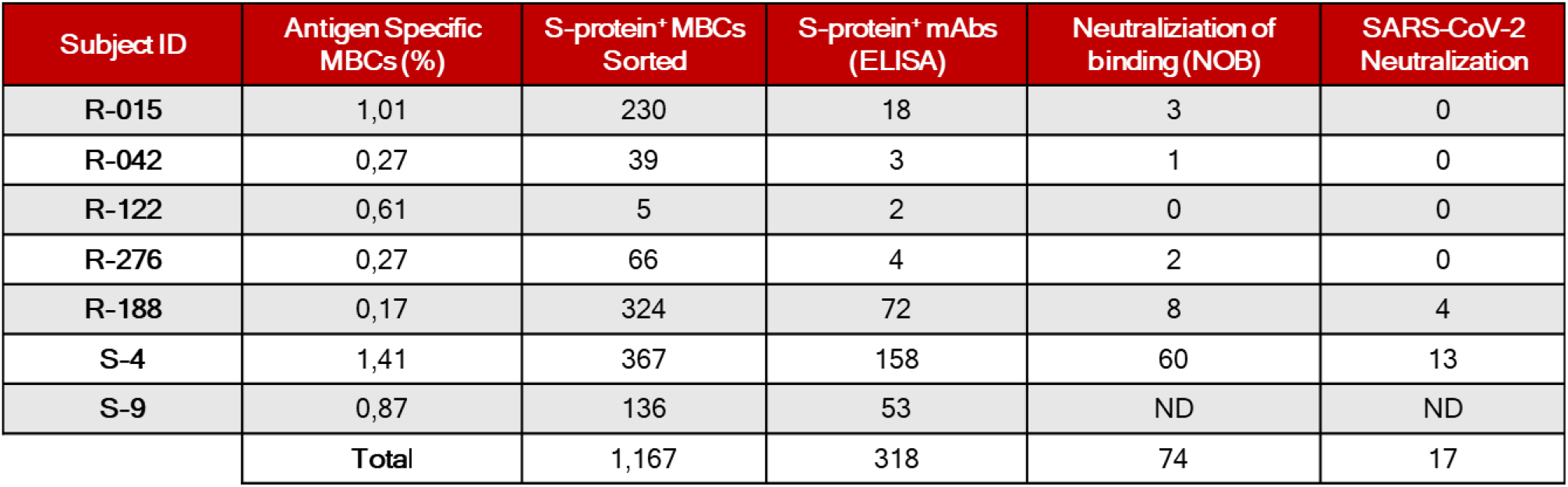
SARS-CoV-2 convalescent donors S-protein specific MBCs analyses. The Table reports the number of S-protein-specific MBCs that were sorted and screened (for binding by ELISA and for functionality by NOB and viral neutralization) for each subject enrolled in this study; ND = Not Done.

### Functional characterization of S-protein specific mAbs against SARS-CoV-2

Of the 318 supernatants containing S-protein specific mAbs, 265 were screened *in vitro* for their ability to block the binding of the S-protein to Vero E6 cell receptors (Table 2 and Fig. 4A) and for their neutralization activity against the SARS-CoV-2 virus (Fig. 5A). In the NOB assay, 74 of the 265 tested (28%) S-protein specific mAbs were able to neutralize the antigen/receptor binding showing a broad array of neutralization potency ranging from 52% to over 90% (Fig. 4B). In the viral neutralization assay, supernatants containing naturally produced IgG or IgA were tested for their ability to protect the layer of Vero E6 cells from the cytopathic effect triggered by SARS-CoV-2 infection (Fig. 5A). When mAbs were able to protect Vero E6 cells from infection, i.e. showing neutralization capacity against SARS-CoV-2, no cytopathic effect was observed (Fig. 5B, bottom-left box). On the contrary, when mAbs were not able to prevent infection, cytopathic effect on Vero E6 was clearly visible (Fig. 5B, bottom-right box). Out of the 265 mAbs tested in this study, a panel of 17 mAbs neutralized the virus and prevented infection of Vero E6 cells (Table 2).

**Fig. 4.**
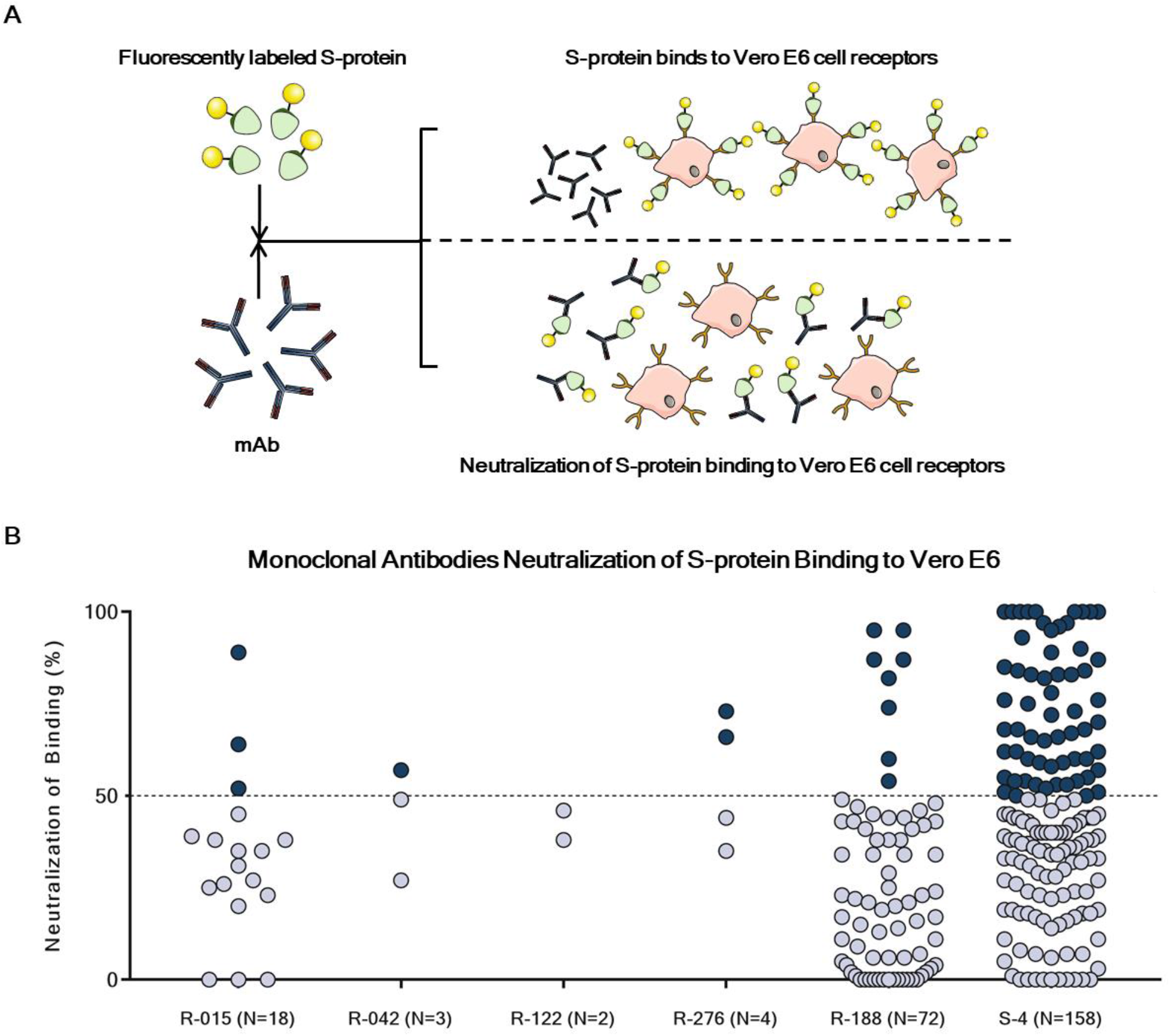
Neutralization of S-protein binding to Vero E6 cell receptors by S-protein specific mAbs. (A) Schematic representation of the neutralization of binding (NOB) assay used to screen isolated S-protein specific mAbs for their ability to abrogate the interaction between SARS-CoV-2 and Vero E6 cell receptors. (B) The graph shows supernatants tested by NOB assay. Threshold of positivity has been set as 50% of binding neutralization (dotted line). Dark blue dots represent mAbs able to neutralize the binding between SARS-CoV-2 and receptors on Vero E6 cells, while light blue dots represent non-neutralizing mAbs. The total number (N) of S-protein specific supernatants screened by NOB assay is shown for each donor.

**Fig. 5.**
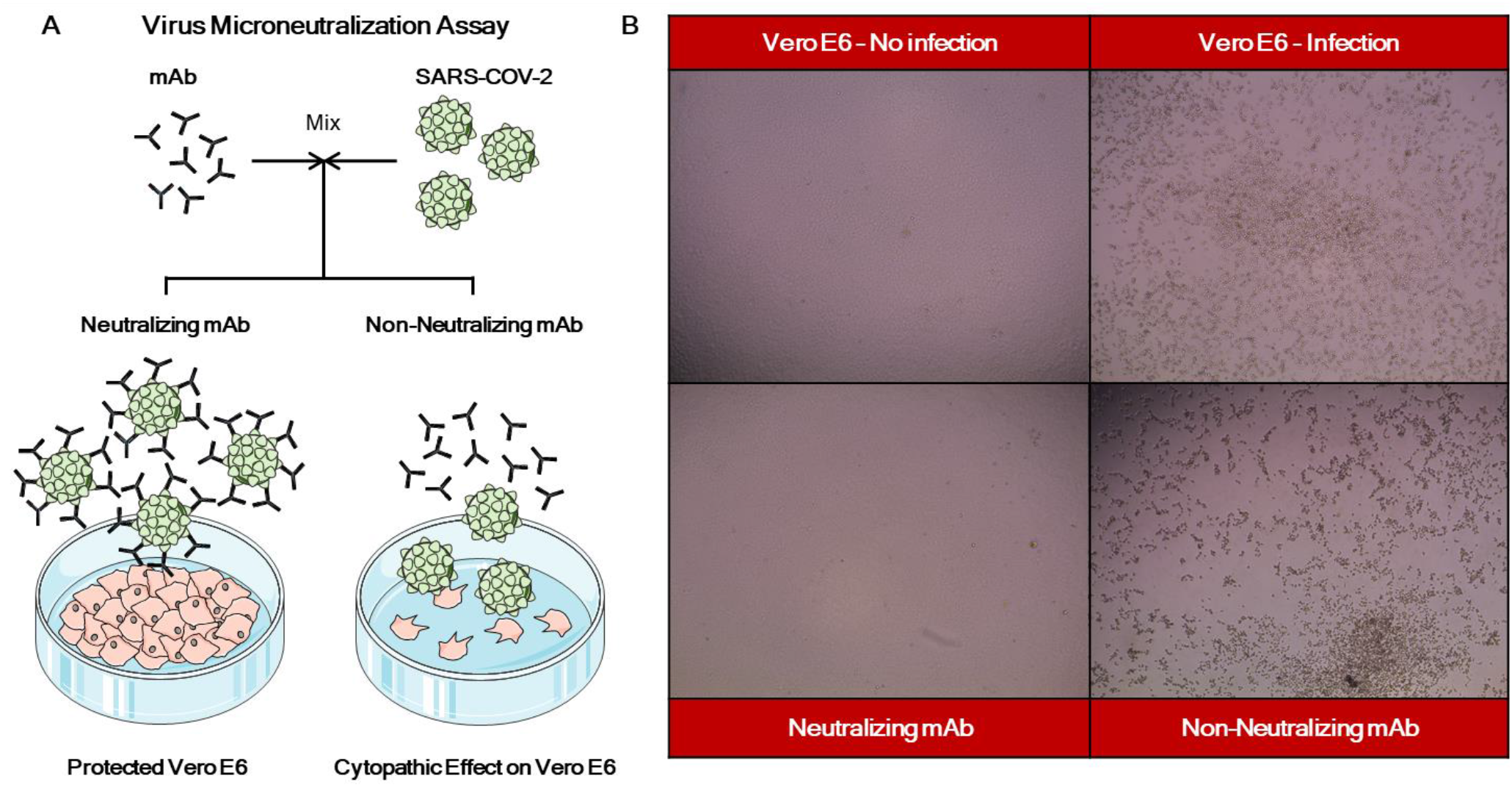
SARS-CoV-2 neutralization assay for S-protein specific mAbs. (A) Schematic representation of the virus neutralization assay used in this study to assess functional activities of S-protein specific mAbs. (B) Representative microscope images showing the cytopathic effect of SARS-CoV-2 or the protective efficacy of the screened supernatants on Vero E6 cells.

## DISCUSSION

Human monoclonal antibodies are an industrially mature technology with more than 50 products already approved in the field of cancer, inflammation and autoimmunity. The well-established safety profile and the large experience for their development, make mAbs ideal candidates for rapid development especially in epidemic and pandemic settings. So far mAbs have rarely been used in the field of infectious diseases, mostly because the large quantities needed for therapy made them not cost effective. However, in recent years, the incredible technological progress to isolate and screen memory B cells made possible to identify highly potent neutralizing mAbs and to further improve their potency of several orders of magnitude through established engineering procedures. This possibility resulted into a decreased amount of antibodies necessary for therapy thus making non-intravenous delivery of potent neutralizing mAbs possible. Several candidates are presently under development in the field of HIV, pandemic influenza, RSV and many other infectious diseases (*11*). Perhaps the most striking demonstration of the power of mAbs for emerging infections came from the Ebola experience. In this case rapidly developed potent mAbs were among the first drugs to be tested in the Ebola outbreak and showed remarkable efficacy in preventing mortality (*2*). Given the remarkable efficacy of this intervention, potent mAbs became the first, and so far the only, drug to be recommended for Ebola by the WHO.

In the case of SARS-CoV-2, where so far we do not have any effective therapeutic nor prophylactic interventions, mAbs have the possibility to become one of the first drugs that can be used for immediate therapy of any patient testing positive for the virus, and even to provide immediate protection from infection in high risk populations. Preliminary evidences showed that plasma from infected people improves the outcome of patients with severe disease, therefore it is highly possible that a therapeutic and/or prophylactic mAb-based intervention can be highly effective (*12*). Furthermore, vaccination strategies inducing neutralizing antibodies have shown to protect non-human primates from infection (*13*). These results further stress the importance of mAbs as a measure to counterattack SARS-CoV-2 infection and to constrain its circulation.

In this work we addressed the question of whether mAbs recognizing SARS-CoV-2 can be recovered from memory B cells of people who survived Covid-19 and whether some of them are able to neutralize the virus. Our data show that SARS-CoV-2 specific mAbs can be successfully isolated from most convalescent donors even if the frequency of S-protein specific memory B cells is highly variable among them. In addition, approximately 28% of isolated mAbs were able to inhibit the binding of the S-protein to the receptor(s) on Vero E6 cells. Finally a fraction of isolated mAbs (N=17) were able to effectively neutralize SARS-CoV-2 with high potency when tested *in vitro*. These data suggest that the method we implemented allows us to successfully retrieve mAbs with potent neutralizing activity against SARS-CoV-2 and we plan to select the most promising candidate(s) for drug development. Lead candidates will be further tested for the ability to generate resistant viruses and for the ability to induce antibody-dependent disease enhancement in appropriate models.

## MATERIALS & METHODS

### Enrollment of SARS-COV-2 convalescent donors and human sample collection

This work results from a collaboration with the National Institute for Infectious Diseases, IRCCS – Lazzaro Spallanzani Rome (IT) and Azienda Ospedaliera Universitaria Senese, Siena (IT) that provided samples from SARS-CoV-2 convalescent donors who gave their written consent. The study was approved by local ethics committees (Parere 18_2020 in Rome and Parere 17065 in Siena) and conducted according to good clinical practice in accordance with the declaration of Helsinki (European Council 2001, US Code of Federal Regulations, ICH 1997). This study was unblinded and not randomized.

### Human peripheral blood mononuclear cells (PBMCs) isolation from SARS-CoV-2 convalescent donors

Peripheral blood mononuclear cells (PBMCs) were isolated from heparin-treated whole blood by density gradient centrifugation (Lympholyte-H; Cedarlane). After separation, PBMC were: i) frozen in liquid nitrogen at concentration of 10 x 10^6^ PBMC/vial using 10% DMSO in heat-inactivated fetal bovine serum (FBS) or ii) resuspended in RPMI 1640 (EuroClone) supplemented with 10% FBS (EuroClone), 2 mmol/L L-glutamine, 2 mmol/L penicillin, and 50 μg/mL streptomycin (EuroClone). Cells were cultured for 18 hour at 37°C with 5% CO_2_. Blood samples were screened for SARS-CoV-2 RNA and for antibodies against HIV, HBV and HCV.

### Expression and purification of SARS-CoV-2 S-protein prefusion trimer

The expression vector coding for prefusion S ectodomain (kind gift of Dr. Jason Mc Lellan) was used to transiently transfect Expi293F cells (Thermo Fisher #A14527) using Expifectamine (Thermo Fisher # A14525). The protein was purified from filtered cell supernatants using NiNTA resin (GE Healtcare #11-0004-58), eluted with 250 mM Imidazole (Sigma Aldrich #56750), dialyzed against PBS, and then stored at 4°C prior to use.

### Single cell sorting of SARS-CoV-2 S-protein^+^ memory B cells

Human peripheral blood mononuclear cells (PBMCs) from SARS-CoV-2 convalescent donors were stained with Live/Dead Fixable Aqua (Invitrogen; Thermo Scientific) in 100 μL final volume diluted 1:500 at room temperature (RT). After 20 min incubation cells were washed with phosphate buffered saline (PBS) and unspecific bindings were saturated with 50 μL of 20% rabbit serum in PBS. Following 20 min incubation at 4°C cells were washed with PBS and stained with SARS-CoV-2 S-protein labeled with Strep-Tactin^®^XT DY-488 (iba-lifesciences cat# 2-1562-050) for 30 min at 4°C. After incubation the following staining mix was used CD19 V421 (BD cat# 562440), IgM PerCP-Cy5.5 (BD cat# 561285), CD27 PE (BD cat# 340425), IgD-A700 (BD cat# 561302), CD3 PE-Cy7 (BioLegend cat# 300420), CD14 PE-Cy7 (BioLegend cat# 301814), CD56 PE-Cy7 (BioLegend cat# 318318) and cells were incubated at 4°C for additional 30 min. Stained MBCs were single cell-sorted with a BD FACSAria III (BD Biosciences) into 384-well plates containing 3T3-CD40L feeder cells and were incubated with IL-2 and IL-21 for 14 days as previously described (*10*).

### ELISA assay with S1 and S2 subunits of SARS-CoV-2 S-protein

The presence of S1- and S2-binding antibodies in culture supernatants of monoclonal S-protein-specific memory B cells was assessed by means of an ELISA assay implemented with the use of a commercial kit (ELISA Starter Accessory Kit, Catalogue No. E101; Bethyl Laboratories, Montgomery, TX, USA). Briefly, 384-well flat-bottom microtiter plates (Nunc MaxiSorp 384-well plates; Sigma-Aldrich) were coated with 25 μl/well of antigen (1:1 mix of S1 and S2 subunits, 1 μg/ml each; The Native Antigen Company, Oxford, United Kingdom) diluted in coating buffer (0.05 M carbonate-bicarbonate solution, pH 9.6), and incubated overnight at 4°C. The plates were then washed three times with 100 μl/well washing buffer (50 mM Tris Buffered Saline (TBS) pH 8.0, 0.05% Tween-20) and saturated with 50 μl/well blocking buffer containing Bovine Serum Albumin (BSA) (50 mM TBS pH 8.0, 1% BSA, 0.05% Tween-20) for 1 hour (h) at 37°C. After further washing, samples diluted 1:5 in blocking buffer were added to the plate. Blocking buffer was used as a blank. After an incubation of 1 h at 37°C, plates were washed and incubated with 25 μl/well secondary antibody (horseradish peroxidase (HRP)-conjugated goat anti-human IgG-Fc Fragment polyclonal antibody, diluted 1:10,000 in blocking buffer, Catalogue No. A80-104P; (Bethyl Laboratories, Montgomery, TX, USA) for 1 h at 37°C. After three washes, 25 μl/well TMB One Component HRP Microwell Substrate (Bethyl Laboratories, Montgomery, TX, USA) was added and incubated 10–15 minutes at RT in the dark. Color development was terminated by addition of 25 μl/well 0.2 M H_2_SO_4_. Absorbance was measured at 450 nm in a Varioskan Lux microplate reader (Thermo Fisher Scientific). The threshold for sample positivity was set at twice the OD of the blank.

### ELISA assay with SARS-CoV-2 S-protein prefusion trimer

ELISA assay was used to detect SARS-CoV-2 S-protein specific mAbs and to screen plasma from SARS-CoV-2 convalescent donors. 384-well plates (Nunc MaxiSorp 384 well plates; Sigma Aldrich) were coated with 3μg/mL of streptavidin diluted in PBS and incubated at RT overnight. Plates were then coated with SARS-CoV-2 S-protein at 3μg/mL and incubated for 1h at room temperature. 50 μL/well of saturation buffer (PBS/BSA 1%) was used to saturate unspecific binding and plates were incubated at 37°C for 1h without CO_2_. Supernatants were diluted 1:5 in PBS/BSA 1%/Tween20 0,05% in 25 μL/well final volume and incubated for 1h at 37°C without CO_2_. 25 μL/well of alkaline phosphatase-conjugated goat anti-human IgG (Sigma-Aldrich) and IgA (Jackson Immuno Research) were used as secondary antibodies. In addition, twelve two-fold serial dilutions of plasma from SARS-CoV-2 infected patients were analyzed in duplicate. Plasma samples were diluted in PBS/BSA 1%/Tween20 0,05% (25 μL/well final volume; Starting Dilution 1:80) and incubated for 1h at 37°C without CO_2_. Next, 25 μL/well of alkaline phosphatase-conjugated goat anti-human IgG (Sigma-Aldrich) was added for 1h at 37°C without CO_2_. Wells were washed three times between each step with PBS/BSA 1%/Tween20 0.05%. PNPP (p-nitrophenyl phosphate) (Thermo Fisher) was used as soluble substrate to detect SARS-CoV-2 S-protein specific monoclonal antibodies and the final reaction was measured by using the Varioskan Lux Reader (Thermo Fisher Scientific) at a wavelength of 405 nm. Samples were considered as positive if optical density at 405 nm (OD_405_) was two times the blank.

### SARS-CoV-2 virus and cell infection

African green monkey kidney cell line Vero E6 cells (American Type Culture Collection [ATCC] #CRL-1586) were cultured in Dulbecco’s Modified Eagle’s Medium (DMEM) - High Glucose (Euroclone, Pero, Italy) supplemented with 2 mM L-Glutamine (Lonza, Milano, Italy), penicillin (100 U/mL) – streptomycin (100 μg/mL) mixture (Lonza, Milano, Italy) and 10% Foetal Bovine Serum (FBS) (Euroclone, Pero, Italy). Cells were maintained at 37°C, in a 5% CO_2_ humidified environment and passaged every 3-4 days.

Wild type SARS CoV-2 2019 (2019-nCoV strain 2019-nCov/Italy-INMI1) virus was purchased from the European Virus Archive goes Global (EVAg, Spallanzani Institute, Rome). For virus propagation, sub-confluent Vero E6 cell monolayers were prepared in T175 flasks (Sarstedt) containing supplemented D-MEM high glucose medium. For titration and neutralization tests of SARS-CoV-2, Vero E6 were seeded in 96-well plates (Sarstedt) at a density of 1,5×10^4^ cells/well the day before the assay.

### Neutralization of Binding (NOB) Assay

To study the binding of the Covid-19 Spike protein to cell-surface receptor(s) we developed an assay to assess recombinant Spike protein specific binding to target cells and neutralization thereof.

To this aim the stabilized Spike protein was coupled to Streptavidin-PE (eBioscience # 12-4317-87, Thermo Fisher) for 30 min at 4°C and then incubated with VERO E6 cells. Binding was assessed by flow cytometry. The stabilized Spike protein bound VERO E6 cells with high affinity (data not shown).

To assess the content of neutralizing antibodies in sera or in B-cell culture supernatants, two microliters of SARS-CoV-2 Spike-Streptavidin-PE at 15-30 μg/ml in PBS-1%FCS were mixed with two microliters of various dilutions of sera or B-cell culture supernatants in U bottom 96-well plates. After incubation at 37°C for 1 hr, 25×10 Vero E6 cells suspended in two microliters of PBS 1% FCS were added and incubated for additional 1 hr at 4°C. Non-bound protein and antibodies were removed and cell-bound PE-fluorescence was analyzed with a FACScantoII flow cytometer (Becton Dickinson). Data were analyzed using the FlowJo data analysis software package (TreeStar, USA). The specific neutralization was calculated as follows: NOB (%) = 1 – (Sample MFI value – background MFI value) / (Negative Control MFI value – background MFI value).

### Viral propagation and titration

The SARS-CoV-2 virus was propagated in Vero E6 cells cultured in DMEM high Glucose supplemented with 2% FBS, 100 U/mL penicillin, 100 μg/mL streptomycin. Cells were seeded at a density of 1×10^6^ cells/mL in T175 flasks and incubated at 37°C, 5% CO_2_ for 18-20 hours. The subconfluent cell monolayer was then washed twice with sterile Dulbecco’s phosphate buffered saline (DPBS). Cells were inoculated with 3.5 ml of the virus properly diluted in DMEM 2% FBS at a multiplicity of infection (MOI) of 0.001, and incubated for 1h at 37°C in a humidified environment with 5% CO_2_. At the end of the incubation, 50 mL of DMEM 2% FBS were added to the flasks. The infected cultures were incubated at 37°C, 5% CO_2_ and monitored daily until approximately 80-90% of the cells exhibited cytopathic effect (CPE). Culture supernatants were then collected, centrifuged at 4°C at 1,600 rpm for 8 minutes to allow removal of cell debris, aliquoted and stored at −80°C as the harvested viral stock. Viral titers were determined in confluent monolayers of Vero E6 cells seeded in 96-well plates using a 50% tissue culture infectious dose assay (TCID50). Cells were infected with serial 1:10 dilutions (from 10-1 to 10-11) of the virus and incubated at 37°C, in a humidified atmosphere with 5% CO_2_. Plates were monitored daily for the presence of SARS-CoV-2 induced CPE for 4 days using an inverted optical microscope. The virus titer was estimated according to Spearman-Karber formula (*14*) and defined as the reciprocal of the highest viral dilution leading to at least 50% CPE in inoculated wells.

### Semi-quantitative live SARS-CoV-2-based neutralization assay

To assess the neutralization titer of anti-SARS-CoV-2 plasma samples from Covid-19 convalescent donors, a semi-quantitative neutralization method was used. Plasma samples were heat-inactivated for 30 minutes at 56°C and 2-fold serially diluted starting from 1:10 to 1:2,560 dilution, then mixed with an equal volume of viral solution containing 100 TCID50 of SARS-CoV-2 diluted in D-MEM high Glucose 2% FBS. After 1 hour incubation at 37°C, 5% CO_2_, 100 μl of the virus-plasma mixture at each dilution was passed to a cell plate containing a sub-confluent Vero E6 cell monolayer. Plates were incubated for 3 days at 37°C in a humidified environment with 5% CO_2_, then checked for development of CPE by means of an inverted optical microscope. The reciprocal of the highest plasma dilution that resulted in more than 50% inhibition of CPE was defined as the neutralization titer.

### Qualitative live SARS-CoV-2-based neutralization assay

The neutralization activity of culture supernatants from monoclonal S-protein-specific memory B cells was evaluated by means of a qualitative live-virus based neutralization assay against a one-point dilution of the samples. Supernatants were mixed in a 1:3 ratio with a SARS-CoV-2 viral solution containing 25 TCID50 of virus (final volume: 30 μl). After 1 hour incubation at 37°C, 5% CO_2_, 25 μl of each virus-supernatant mixture was added to the wells of a 96-well plate containing a sub-confluent Vero E6 cell monolayer. Following a 2-hour incubation at 37°C, the virus-serum mixture was removed and 100 μl of DMEM 2% FBS were added to each well. Plates were incubated for 3 days at 37°C in a humidified environment with 5% CO_2_, then examined for CPE by means of an inverted optical microscope. Absence or presence of CPE was defined by comparison of each well with the positive control (plasma sample showing high neutralizing activity of SARS-CoV-2 in infected Vero E6 cells) and negative control (human serum sample negative for SARS-CoV-2 in ELISA and neutralization assays).

## Author contribution

Emanuele Andreano, Ida Paciello isolated single memory B cells and identified S-protein specific mAbs (cell sorting and ELISA).

Piero Pileri, Noemi Manganaro, Elisa Pantano performed NOB assay and produced recombinant spike protein.

Giulia Piccini, Alessandro Manenti and Emanuele Montomoli performed ELISA and viral neutralization assay.

Marco Troisi, Fabiola Vacca, Concetta De Santi, Dario Cardamone, Anna Kabanova contributed to the characterization of positive memory B cells.

Emanuele Nicastri, Chiara Agrati, Concetta Castilletti, Francesca Montagnani, Arianna Emiliozzi, Massimiliano Fabbiani, Maria Rosaria Capobianchi enrolled patients and isolated PBMCs.

Claudia Sala, Giuseppe Ippolito, Rino Rappuoli coordinated the project.

## ACKNOWLEDGMENTS

We wish to thank Fondazione Toscana Life Sciences in the persons of Dr. Fabrizio Landi and Dr. Andrea Paolini and the whole Administration for their incredible help and support. In particular we would like to thank Mr. Francesco Senatore, Mrs. Laura Canavacci and Mrs. Cinzia Giordano for their support in preparing all the documents needed for the ethical approval of the clinical studies carried out within this project.

This work was possible thanks to the technology established in our European Research Council (ERC) funded laboratory through the ERC Advanced grant vAMRes which allowed us to isolate mAbs from vaccinated and/or convalescent patients to tackle the global threat posed by antimicrobial resistance.

We wish to thank the National Institute for Infectious Diseases, IRCCS, Lazzaro Spallanzani Rome (IT) and the Azienda Ospedaliera Universitaria Senese, Siena (IT), for providing blood samples from Covid-19 convalescent donors under studies approved by local ethic committees. We also wish to thank all the nursing staff who chose to cooperate for blood withdrawal and all the donors who decided to participate in this study.

We would like to thank the whole GSK Vaccines Pre-clinical Evidence Generation and Assay – Immunolgy function led by Dr. Oretta Finco for their availability and support as well as Mrs. Simona Tavarini, Mrs. Chiara Sammicheli, Dr. Monia Bardelli, Dr. Michela Brazzoli, Dr. Elisabetta Frigimelica, Dr. Erica Borgogni and Dr. Elisa Faenzi for sharing their expertise, extreme availability and technical support. We would also like to thank Dr. Mariagrazia Pizza and Dr. Simone Pecetta for initial insightful advice and discussions on this project.

We would like to thank Dr. Jason McLellan and his team for generously providing the SARS-CoV-2 S-protein stabilized in its prefusion conformation used in this study. Furthermore, we would like to thank Dr. Daniel Wrapp and Dr. Nianshuang Wang for the precious information and suggestions.

We gratefully acknowledge the Collaborators Members of INMI COVID-19 study group: Maria Alessandra Abbonizio, Amina Abdeddaim, Fabrizio Albarello, Gioia Amadei, Alessandra Amendola, Mario Antonini, Tommaso Ascoli Bartoli, Francesco Baldini, Raffaella Barbaro, Bardhi Dorian, Barbara Bartolini, Rita Bellagamba, Martina Benigni, Nazario Bevilacqua, Gianlugi Biava, Michele Bibas, Licia Bordi, Veronica Bordoni, Evangelo Boumis, Marta Branca, Donatella Busso, Marta Camici, Paolo Campioni, Alessandro Capone, Cinzia Caporale, Emanuela Caraffa, Ilaria Caravella, Fabrizio Carletti, Adriana Cataldo, Stefano Cerilli, Carlotta Cerva, Roberta Chiappini, Pierangelo Chinello, Carmine Ciaralli, Stefania Cicalini, Francesca Colavita, Angela Corpolongo, Massimo Cristofaro, Salvatore Curiale, Alessandra D’Abramo, Cristina Dantimi, Alessia De Angelis, Giada De Angelis, Maria Grazia De Palo, Federico De Zottis, Virginia Di Bari, Rachele Di Lorenzo, Federica Di Stefano, Gianpiero D’ûffizi, Davide Donno, Francesca Faraglia, Federica Ferraro, Lorena Fiorentini, Andrea Frustaci, Matteo Fusetti, Vincenzo Galati, Roberta Gagliardini, Paola Gallì, Gabriele Garotto, Saba Gebremeskel Tekle, Maria Letizia Giancola, Filippo Giansante, Emanuela Giombini, Guido Granata, Maria Cristina Greci, Elisabetta Grilli, Susanna Grisetti, Gina Gualano, Fabio Iacomi, Giuseppina Iannicelli, Eleonora Lalle, Simone Lanini, Daniele Lapa, Luciana Lepore, Raffaella Libertone, Raffaella Lionetti, Giuseppina Liuzzi, Laura Loiacono, Andrea Lucia, Franco Lufrani, Manuela Macchione, Gaetano Maffongelli, Alessandra Marani, Luisa Marchioni, Raffaella Marconi, Andrea Mariano, Maria Cristina Marini, Micaela Maritti, Alessandra Mastrobattista, Giulia Matusali, Valentina Mazzotta, Paola Mencarini, Silvia Meschi, Francesco Messina, Annalisa Mondi, Marzia Montalbano, Chiara Montaldo, Silvia Mosti, Silvia Murachelli, Maria Musso, Pasquale Noto, Roberto Noto, Alessandra Oliva, Sandrine Ottou, Claudia Palazzolo, Emanuele Pallini, Fabrizio Palmieri, Carlo Pareo, Virgilio Passeri, Federico Pelliccioni, Antonella Petrecchia, Ada Petrone, Nicola Petrosillo, Elisa Pianura, Carmela Pinnetti, Maria Pisciotta, Silvia Pittalis, Agostina Pontarelli, Costanza Proietti, Vincenzo Puro, Paolo Migliorisi Ramazzini, Alessia Rianda, Gabriele Rinonapoli, Silvia Rosati, Martina Rueca, Alessandra Sacchi, Alessandro Sampaolesi, Francesco Sanasi, Carmen Santagata, Alessandra Scarabello, Silvana Scarcia, Vincenzo Schininà, Paola Scognamiglio, Laura Scorzolini, Giulia Stazi, Fabrizio Taglietti, Chiara Taibi, Roberto Tonnarini, Simone Topino, Francesco Vaia, Francesco Vairo, Maria Beatrice Valli, Alessandra Vergori, Laura Vincenzi, Ubaldo Visco-Comandini, Pietro Vittozzi, Mauro Zaccarelli.

We would like to thank all the other Members of the Covid-19 UNIT and of the Covid-19 TEAM, AOUS (Azienda Ospedaliera Universitaria Senese, Siena, Italy): Giovanni Antonelli, Tommaso Bacconi, Riccardo Barbati, Claudia Basagni, David Bennett, Francesco Bianchi, Cesare Biuzzi, Francesco Bova, Carla Caffarelli, Paolo Cameli, Chiara Cassol, Elena Ceccarelli, Raffaella Corbisiero, Lucia Cubattoli, Maria Grazia Cusi, Vincenzo De Franco, Antonio De Siero, Emanuele Detti, Giovanni Donati, Lediona Dyrmo, Camilla Fabbri, Federico Franchi, Bruno Frediani, Severino Gallo, Benedetta Galgani, Rodolfo Gentilini, Elisa Giacomin, Roberto Gusinu, Domenico Iemma, Anna Incisore, Alessandro Lanari, Nicola Lanzarone, Andrea Lapi, Stefano Lunghetti, Marco Mautone, Riccardo Marcucci, Adriana Chiara Marinetti, Daniele Marri, Maria Laura Marzi, Melissa Masini, Egidio Mastrocinque, Chiara Mattaliano, Maria Antonietta Mazzei, Maria Mencarelli, Fabrizio Mezzasalma, Lucia Migliorini, Rosita Morelli, Matteo Nardi, Lorenzo Paglicci, Francesco Palilla, Paolo Petrioli, Francesco Pippi, Daria Pizzirani, Aristotele Porceddu, Barbara Porchia, Salvatore Quarta, Barbara Rossetti, Irene Sellerio, Anna Sansoni, Amato Santoro, Sabino Scolletta, Irene Sellerio, Giulia Stella, Pitinca Tomai, Serafina Valente, Luca Volterrani, Giacomo Zanelli, and the Covid-19 nursing staff.

This publication was supported by funds from the “Centro Regionale Medicina di Precisione” and by all the people who answered the call to fight with us the battle against SARS-CoV-2 with their kind donations on the platform ForFunding (https://www.forfunding.intesasanpaolo.com/DonationPlatform-ISP/nav/progetto/id/3380).

This publication was supported by the European Virus Archive goes Global (EVAg) project, which has received funding from the European Union’s Horizon 2020 research and innovation programme under grant agreement No 653316.

